# Antisense RNA C9orf72 Hexanucleotide Repeat Associated With Amyotrophic Lateral Sclerosis and Frontotemporal Dementia Forms A Triplex-Like Structure and Binds Small Synthetic Ligand

**DOI:** 10.1101/2023.10.10.561654

**Authors:** Leszek Błaszczyk, Marcin Ryczek, Bimolendu Das, Martyna Mateja-Pluta, Magdalena Bejger, Kazuhiko Nakatani, Agnieszka Kiliszek

**Affiliations:** Institute of Bioorganic Chemistry, Polish Academy of Sciences, Z. Noskowskiego 12/14, 61-704 Poznań, Poland; Department of Regulatory Bioorganic Chemistry, SANKEN (The Institute of Scientific and Industrial Research), Osaka University, 8-1 Mihogaoka, Ibaraki, Osaka 567-0047, Japan

**Keywords:** RNA structure, RNA-ligand interactions, RNA triplex, hexanucleotide repeat expansion, ALS/FTD disorder

## Abstract

The abnormal expansion of GGGGCC/CCCCGG hexanucleotide repeats (HR) in C9orf72 is associated with familial amyotrophic lateral sclerosis (ALS) and frontotemporal dementia (FTD). Structural polymorphisms of HR result in the multifactorial pathomechanism of ALS/FTD. Consequently, many ongoing studies are focused at developing therapies targeting pathogenic HR RNA. One of them involves small molecules blocking the sequestration of important proteins, preventing the formation of toxic nuclear foci. However, rational design of potential therapeutics is hindered by limited number of structural studies of RNA-ligand complexes. We determined the crystal structure of antisense HR RNA in complex with ANP77 ligand and in the free form. HR RNA folds into a triplex structure composed of four RNA chains. ANP77 interacted with two neighboring single-stranded cytosines to form pseudo-canonical base pairs by adopting sandwich-like conformation and adjusting the position of its naphthyridine units to the helical twist of the RNA. In the unliganded structure, the cytosines formed a peculiar triplex i-motif, assembled by trans C•C+ pair and a third cytosine located at the Hoogsteen edge of the C•C+ pair. These results extend our knowledge of the structural polymorphisms of HR and can be used for the rational design of small molecules targeting disease-related RNAs.

## INTRODUCTION

Amyotrophic Lateral Sclerosis and Frontotemporal Degeneration are fatal neurodegenerative repeat expansion disorders that affect the motor neurons in the brain and spinal cord. The progressive degeneration of nerve cells leads to changes in behavior, dysphagia, dysarthria, respiratory failure, and consequently, death.^1^ Currently, only symptomatic treatments are available to alleviate disease progression. The most frequent cause of ALS/FTD is a mutation in the C9orf72 gene, which encodes a protein involved in the autophagy-lysosome pathway.^2^ The promoter region of C9orf72 caries microsatellite sequences consisting of hexanucleotide repeats (HR) 5’-GGGGCC3’/5’-GGCCCC-3’, which can undergo abnormal expansion. Healthy individuals possess up to 20 repeats, whereas the mutated form of the gene contains several thousand HR units.^2, 3^ The presence of expanded repeats has a negative effect on DNA and RNA functions, resulting in the complex pathomechanism of ALS/FTD. One of these pathways involves transcriptional gene silencing and activation of the DNA damage response through the formation of DNA-RNA hybrids (R-loops).^2, 4^ On the RNA level, bidirectional transcription produces sense and antisense transcripts containing complementary regions of repeated stretches of GGGGCC (G4C2) and GGCCCC (G2C4) that form stable secondary and tertiary structures.^4, 5^ As a consequence, mutated transcripts gain the ability to sequester important proteins and form RNA foci that accumulate in the nucleus.^6^ Expanded repeats can also trigger repeat-associated non-ATG (RAN) translation, resulting in toxic cellular polydipeptide aggregates.^7^

Sense and antisense transcripts containing HR fold into tertiary motifs and exhibit structural polymorphisms. Sense G4C2 RNA is rich in guanine residues.

Thus, in the presence of potassium ions, interor intramolecular G-quadraplexes can be formed and assembled into G-wire or gel-like phases.^4, 5, 8, 9^ G4C2 RNA repeats also fold into hairpin structures, which dominate in the absence of K+ ions or at low refolding temperature.^4, 5, 9^ The antisense G2C4 repeats are cytosine rich and can fold into hairpin structures with long helical stems. Alternatively, cytosines can undergo protonation to form imotifs or triplexes.^4, 5, 10^ So far, only one crystal structure of a G2C4 repeat has been reported.^11^ It folds into a duplex, representing part of the hairpin stem, consisting of G-C pairs interposed by non-canonical C-C pairs. The 3D structures of G2C4 repeats with protonated cytosine residues are currently unknown.

The structural polymorphism of the expanded HR has implications for the development of drugs against ALS/FTD. One proposed therapy involves small molecules that can interact with expanded HR and block its protein-binding sites, preventing the formation of toxic foci and inhibiting RAN translation.^9^ Most investigations have focused on targeting G4C2 repeats with small molecules using library screening, whereas the druggability of G2C4 repeats has not been explored.^9, 12, 13^ To be effective, targeted therapies require the development of ligands exhibiting high structural specificity against target RNA.^13, 14^ The rational design of lead compounds and potential therapeutics can be greatly improved by threedimensional structures of RNA in a ligand-bound form. However, this knowledge is limited with regards to disease-related repeat sequences.^14, 15^

Here, we present the crystal structure of G2C4 RNA in complex with the synthetic molecule ANP77. RNA folds into a triplex-like structure involving protonated cytosine residues, whereas ANP77 interacts directly with RNA via pseudo-canonical base pairs. These crystallographic models are rare examples demonstrating unexplored potential of cytosine-rich sequences to form complex RNA structures and provide sophisticated crystallographic templates for structure-guided approaches in the development of lead compounds for drug discovery against ALS/FTD.

## RESULTS AND DISCUSSION

### The overall structure of the G2C4-ANP77 complex

The ANP77 ligand was designed to selectively bind to the internal loop of the Y/CC motif in RNA (Y denotes pyrimidine).^16, 17^ It is composed of two 2-amino-1,8-naphthyridine units connected by a short aliphatic linker (Figure 1A-C). Each naphthyridine unit (NU) served as a hydrogen bonding partner for interactions with cytosine residues. At a weakly acidic or neutral pH, protonation occurred at one of the NUs, whereas protonation of the second unit required a lower pH (Figure 1A and 1B).^17^

**Figure 1.**
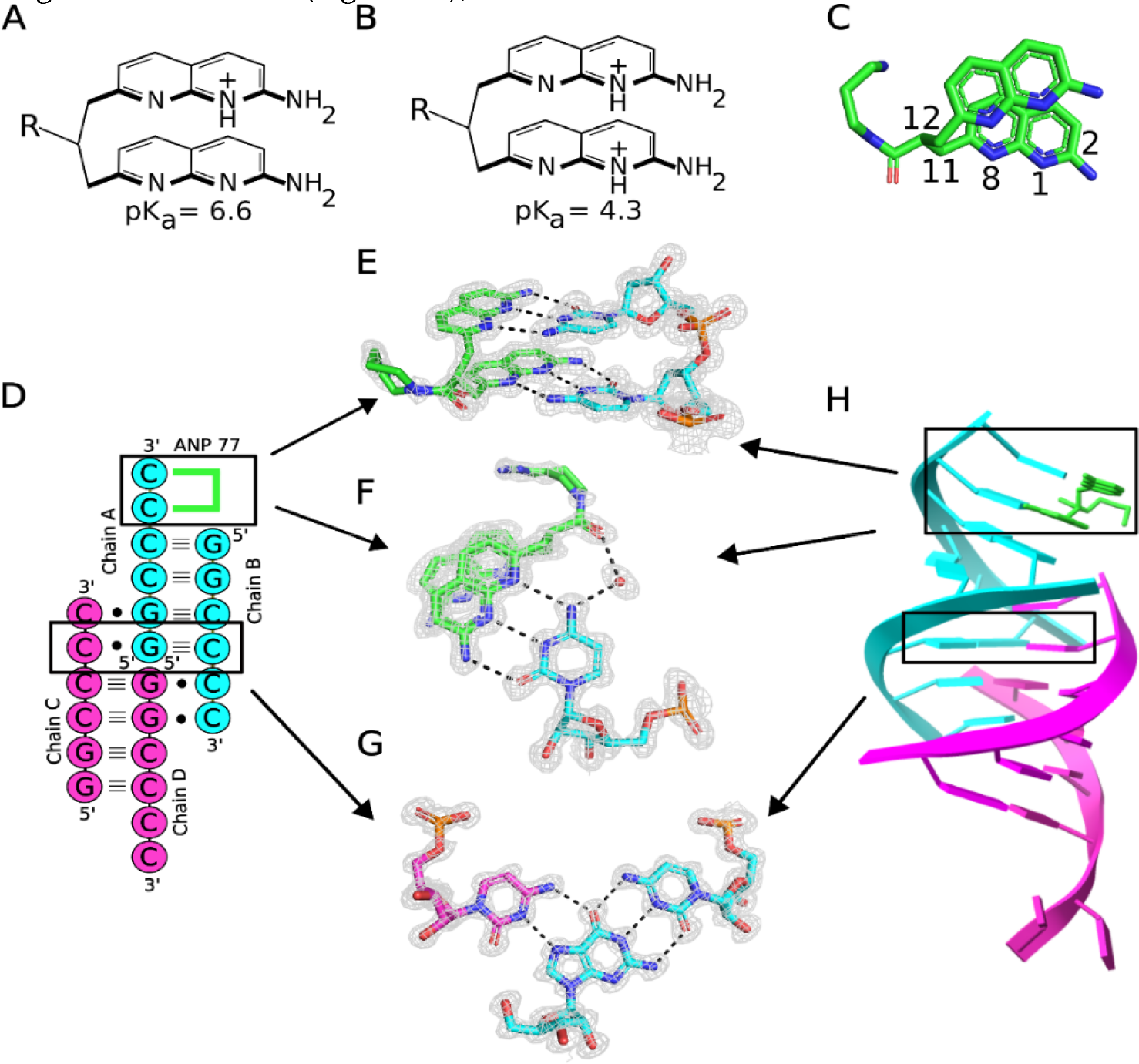
Interactions of ANP77 with G2C4 RNA. (A-B) chemical structure of the ANP77 molecule: (A) in single and (B) doubleprotonated state. (C) stacked conformation of ANP77 observed in the crystal structure. Selected atoms are indicated (see text for details) (D) secondary structure of G2C4-ANP77 complex. Green lines represent ANP77 ligand. (E-F) Pseudo-canonical basepairs formed between ANP77 (green) and cytosine residues from chain A (cyan). (G) Base triple between cytosine from chain C (purple), guanosine from chain A (cyan), and cytosine from chain B. (H) Crystal model of RNA tetramer (chain A and B in cyan; chain C and D in pink) with bound ligand molecule (green sticks). The 2Fo–Fc electron density map (gray) is contoured at the 1σ level. The H-bonds are represented by black dashed lines.

The three-dimensional model of the G2C4-ANP77 complex exhibited a complicated triplex-like fold structure composed of four chains of G2C4 RNA (Protein Data Bank code 8QMH) (Table S1). Chains A and B assembled into duplex AB, whereas chains C and D assembled into duplex CD (Figure 1D). In each duplex, four canonical G-C pairs were present as well as two singlestranded, overhanging cytosines at each 3’ end (Figure 1D). In the crystal lattice, duplexes AB and CD interacted with each other by docking 3’ overhanging cytosines from one end of the duplex into each other’s major grooves (Figure 1D). Cytosine from chain B interacted with duplex CD, whereas cytosine from chain C interacted with duplex AB. Consequently, chains A and D were arranged headto-head, forming a pseudo-continuous strand. Overall, the RNA folded into parallel triplex-like motif flanked by double-stranded duplexes and terminated by singlestranded cytosines (Figure 1D and 1H). The triplex motif was composed of four C+•G-C pairs representing standard major groove base triples (C+ denotes protonated cytosine).^18, 19^ G and C formed canonical Watson-Crick pairs, whereas C+ was located in the major groove and interacted with the Hoogsteen edge of guanosine from the G-C pair. The C+•G interaction included two H-bonds: one between the exo-amino group of C and the O6 carbonyl of G, and the second bond between the N3 amino group of C and the N7 imino atom of G (Figure 1G). The latter interaction required the protonation of cytosine, resulting in the conversion of the N3 imino atom of C+ to an amino function.

In previous studies, G2C4 RNA was crystallized as a duplex.^11^ The cytosines were engaged in canonical pairing with guanosines and in the formation of noncanonical C-C pairs. Although cytosines can interact with each other to form H-bonds, crystallographic studies of C-rich sequences, such as CCG or CCUG repeats, have indicated that the system tends to minimize the number of C-C pairs while simultaneously maximizing the number of G-C pairs.^11, 20^ This was achieved by strand slippage, resulting in 5’ or 3’ overhanging residues or the formation of other types of non-canonical pairs i.e. C-U instead of C-C.^20^ Our study confirmed this tendency. The G2C4 RNA formed slippery duplexes with two 3’ overhanging cytosines. In contrast to other studies, the terminal cytosines were ordered, and some were also engaged in the formation of a triplex structure.

### Pseudocanonical ANP77–cytosine pairs

In the G2C4-ANP77 structure, one ligand molecule was present. It interacted with the two overhanging cytosines of chain A, forming two pseudo-canonical base pairs (Figure 1D and 1H). The NUs of ANP77 mimicked the nitrogen bases of nucleotides, forming three H-bonds with the functional groups of the overhanging cytosines from chain A at their Watson-Crick edges (Figure 1E and 1F). A hydrogen bond was observed between the 2-amino group of ANP77 and the O2 carbonyl atom of cytosine, a second bond between the N8 imino group of the ligand and the exo-amino group of C, and a third interaction between the N1 atom of naphthyridine and the N3 function of cytosine. This H-bonding implies the protonation of one of the nitrogen atoms. Despite the high quality of the electron density map, the localization of the protons within the two pseudo-pairs was ambiguous (Figure 1F). One proton was most likely located at one of the NUs because, at neutral pH, ANP77 is single-charged. The second proton can be attributed to either cytosine or second NU. The second pair of overhanging cytosines from chain D was ordered and interacted with symmetry-related RNA and the 2-exo amino group of the ligand (Figure S1).

### Conformation of ANP77 ligand

In the presented model, ANP77 exhibited a sandwich-like conformation, with the NUs shifting in relation to each other, resulting in limited stacking interactions between the aromatic moieties of the ligand (Figure 1C and 1E). One of the NUs overlapped extensively with the neighboring 1G from chain B (Figure 1H), while the second unit did not form stacking interactions, despite its close proximity to the G-C pair of symmetryrelated RNA molecules.

The aminopropyl carboxamide side chain of ANP77 was located in the major groove of RNA. Its amino-terminal region was disordered, indicating conformational freedom. Only the carbonyl group of the amide bond was modeled; however, it was in a double conformation. Depending on the position of the carbonyl group, it interacted with water molecules from the Hoogsteen edge of 5C involved in pseudo-pairing or with water molecules bridging the Hoogsteen edge of 4C from chain A (Figure 1F).

The crystallographic model of the G2C4-ANP77 complex confirmed that the ligand was capable of interacting directly with two consecutive cytosines. Through the formation of pseudo-base pairs, it merged into an RNA helix, extending the length of the double-stranded region (Figure 1H). This indicates that ANP77 fits into the structure not only in terms of H-bonding, but also in terms of the shape of the RNA helix. The geometry of the linker allowed the positioning of its aromatic rings in a similar manner as the nitrogen bases arranged in a nucleic acid helix. The NUs stacked one above another and were approximately 3 Å apart, reflecting the value of rise parameter in A-RNA form. Moreover, they were twisted to each other by approximately 25°, and the degree of twist could easily be adjusted by rotation around the C11C12 bond (Figure 1C). The length of the propane linker seemed to be properly selected, as it allowed the binding of only two consecutive cytosines. Adding more carbon atoms would increase the flexibility and conformational freedom of the ligand, which could have a negative impact on ANP77 selectivity.

### How to improve the properties of the ANP77 ligand Our model provide the opportunity to rationally

design modifications of ANP77 to improve its interaction with RNA in terms of the specificity, stability, and structural features of the target sequence. One modification would be to induce the protonation of both NUs because, at a neutral pH, only one of the nitrogen atoms (N1 or N1’) possesses proton.^17^ In the crystal structure, it acted as a hydrogen-bond donor for cytosine’s N3 imino group. In turn, a maximum of three H-bonds were formed between naphthyridine and cytosine. Securing the protonation of both units could facilitate tight binding and increase the ligand affinity for the target RNA. The second modification could involve redesigning the side chain of the ligand, which is located close to the major groove of the helix. The electrostatic potential generated by exo-amino groups of cytosines was predominantly positive, and most of exo-amino groups interacted with water molecules, indicating their binding potential. Replacing the protonated amine located at the end of the side chain with a more negatively charged group could be more beneficial. One possible example is the hydroxyl group, which has an electronegative character and can serve as an acceptor of the H-bond.

### Unliganded structure of G2C4 RNA

To determine whether the unusual RNA folding observed in the liganded structure was induced by the binding of ANP77, we determined the crystal structure of G2C4 in its free form (Protein Data Bank code 8QMI) (Table S1). The overall assembly of the triplex structure remained the same. The two terminal cytosines of chain A, which were bound to ANP77 in the liganded structure, engaged in higher-order interactions (Figure 2A). They formed peculiar C•C+•C base triples assembled by symmetry-related residues (5C of chain A, 6C of symmetryrelated chain A, and 6C of symmetry-related chain D) (Figures 2A and 2C). The base triples consisted of a trans C•C+ pair commonly found in i-motif structures. The Nglycosidic bonds were oriented in trans relative to each other and one of the cytosines (the N3 atom) was protonated. The C•C+ pair formed three hydrogen bonds via Watson-Crick edges. The third C from the triples was located in the Hoogsteen edge of one of the cytosine residues from the i-motif pair. It interacted with two Hbonds between the wedged cytosine and the cytosine from the C•C+ pair (Figure 2C).

**Figure 2.**
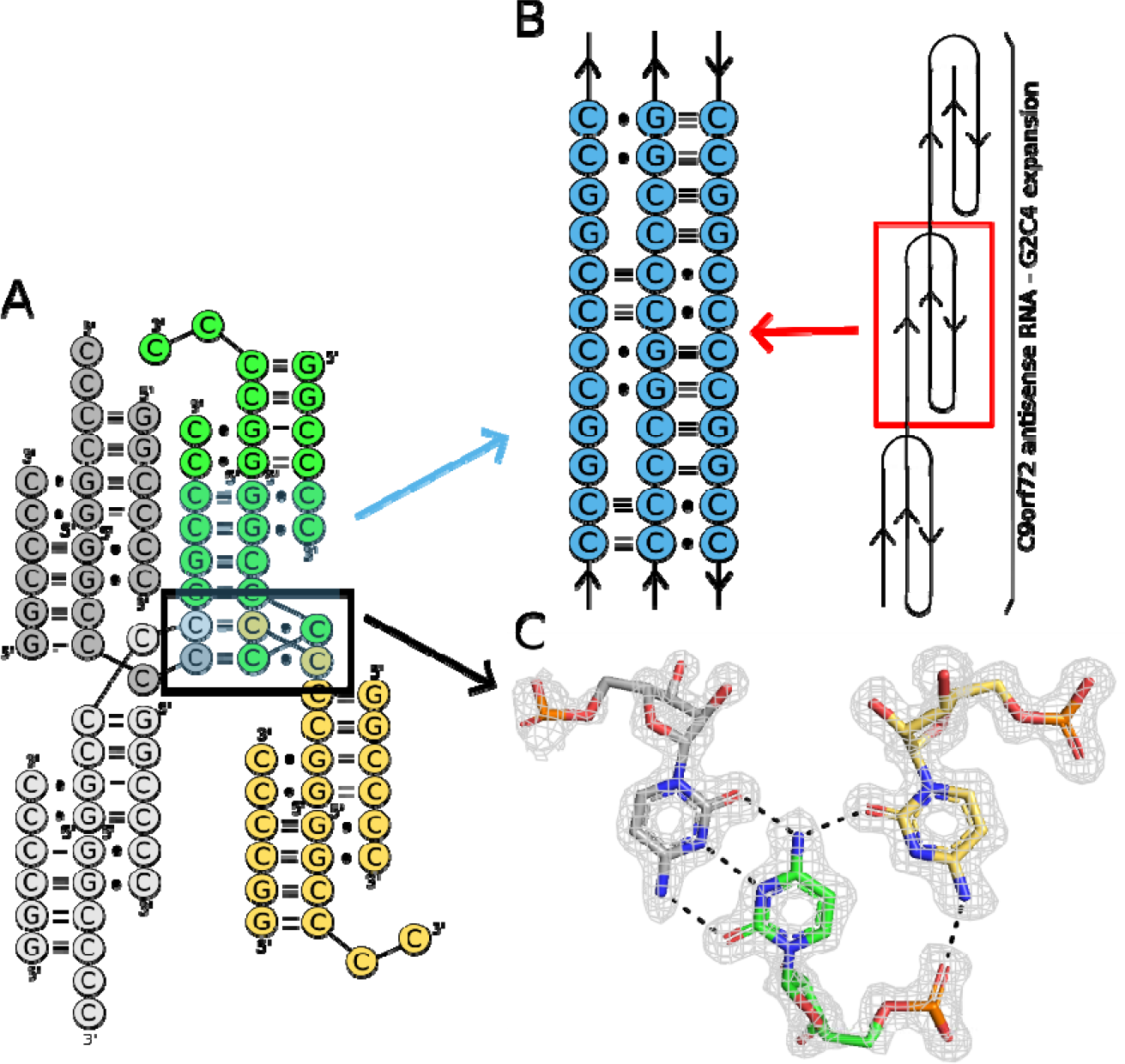
Higher-order interactions of unliganded RNA containing G2C4 repeats. (A) Cytosine residues of three symmetryrelated molecules (grey, yellow and green) form two unique C•C+•C base triples (black box). (B) Proposed model of triplex structure formed by expanded G2C4 repeats (see text for details). The 2Fo–Fc electron density map (grey) is contoured at the 1σ level. The H-bonds are represented by black dashed lines. (C) The C•C+•C triple is assembled by the trans C•C+ pair (green and grey sticks) found in the i-motif structure and cytosine (yellow sticks) located in the Hoogsteen edge moiety.

The unique triplex-i-motif observed in ligand-free G2C4 RNA has not yet been described. This emphasizes the potential of cytosines to assist in the formation of higher-order RNA structures, presenting an example of diverse RNA folding pathways. In our opinion, the structural richness of G2C4 RNA could be implemented in structural biology, that is, in the folding of RNA nanoparticles, the assembly of RNA into multimers for CryoEM measurements, or the enhancement of crystal lattice formation during crystallization. In the context of expanded G2C4 repeats, the obtained results indicated that the G2C4 RNA antisense strand can fold not only into simple hairpins, but also into diverse three-dimensional structures (triplexes or i-motifs). Based on our data, we propose a model in which the long tracts of G2C4 repeats form long triplexes consisting of three different base triples: C•C+•C, C+•G-C, and G•C-G (Figure 2B). This structural arrangement allows cytosines to engage in more favorable H-bond interactions than those observed in the crystal structure of the G2C4 duplex. Triplexes are involved in the folding of RNAs into complex threedimensional architectures, and they are crucial for the biological activity of RNA, such as telomere synthesis, ribosomal frame-shifting, regulation of gene expression through metabolite sensing, and the protection of RNA from degradation.^21, 22^ The triple RNA helix formed by G2C4 repeats can be another example of a triplex that plays an important role in cells, exemplifying the pathological role of HRs.

### Biochemical analysis of G2C4-ANP77 complex Differential scanning calorimetry (DSC) was used

to assess the thermodynamics of ligand binding and folding of G2C4 RNA. Similarly to crystallization experiments, we performed measurements at pH 6.0. Two peaks were observed for the unliganded RNA, indicating melting of the RNA structure: Tm1 at 37.4 °C and Tm2 at 57.2 °C (Figure 3A). In the presence of ANP77, Tm1 was shifted to 42.0 °C, while Tm2 remained unchanged (57.9 °C) (Figure 3B and Table S2). Considering the importance of pH for the protonation of cytosines in terms of the folding of G2C4 RNA into a triplex structure and ligand binding, we performed DSC measurements at pH 7.0 and 5.3. At pH 7.0, for unliganded G2C4, only a single peak was detected at 55.9 °C (1.3 °C lower than Tm2 at pH 6.0) (Figure 3A). The addition of the ligand did not alter the DSC profile (single peak at 54.9 °C) (Figure 3B and Table S2). At pH 5.3, one peak was also detected, but at higher temperatures, either for RNA alone (Tm = 60.0 °C) and the RNAligand complex (Tm = 61.2 °C). However, the height and profile of the peak at pH 5.3 was different from those at pH 6.0 and 7.0.

**Figure 3.**
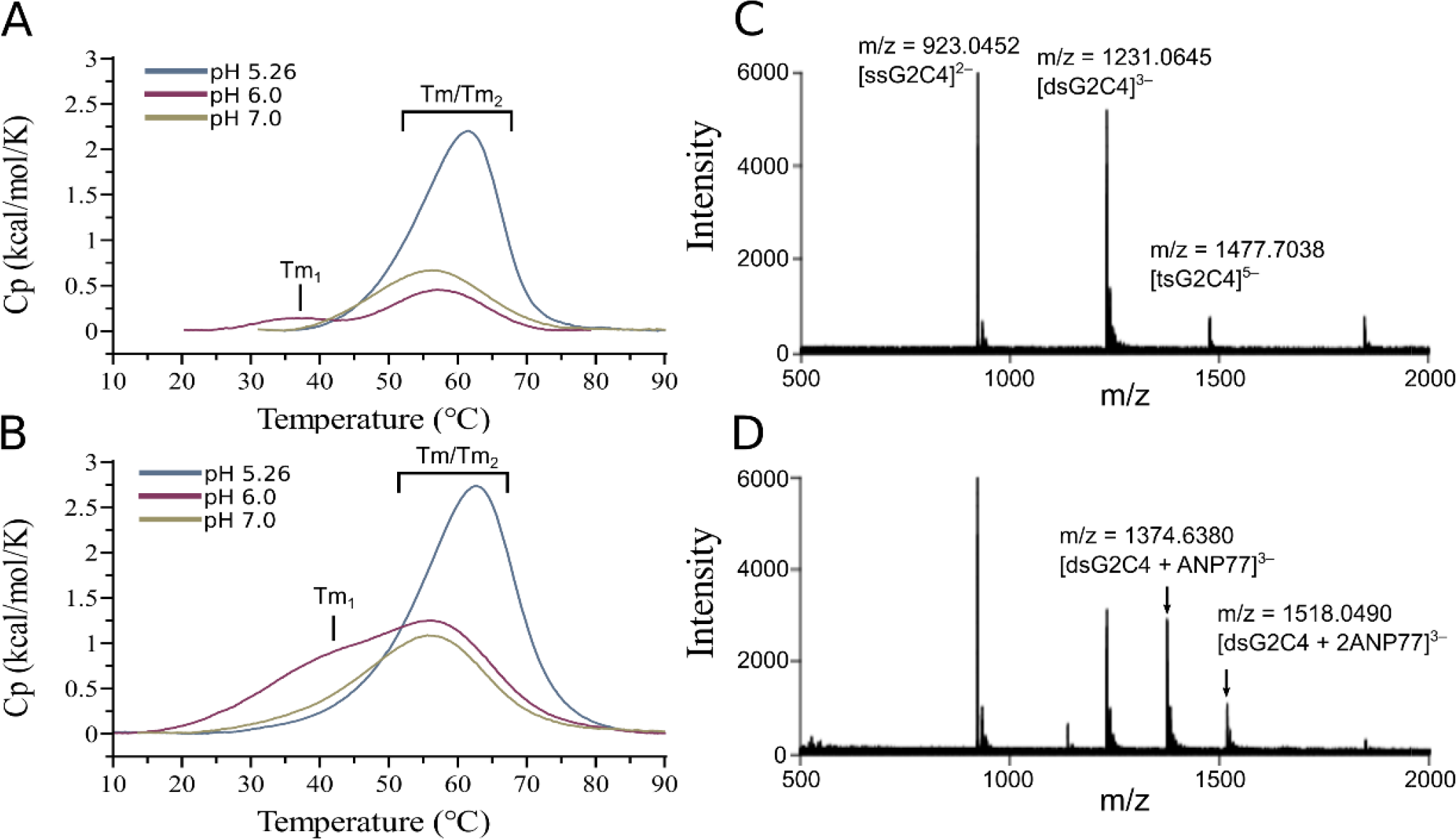
Bio-physical evaluation of interactions between RNA and ANP77 ligand. (A-B) Differential scanning calorimetry (DSC) spectra measured for (A) RNA oligomer and (B) for RNA-ligand complex in different pH conditions. (C-D) Cold-spray ionization time-of-flight mass spectrometry (CSI-TOF-MS) of G2C4 RNA (20 mM) in the (C) absence and (D) presence of ANP77 (50 mM). Asterisk (*) represents an ion corresponding to the 2– ion of dsG2C4 overlapping with the 1– ion of ssG2C4. ssG2C4, dsG2C4, and tsG2C4 denote single-stranded, double-stranded, and tetra-stranded G2C4 RNA, respectively.

Mass spectra indicated the presence of double- and tetra-stranded form of unliganded G2C4 RNA and only a double-stranded form of the RNA-ligand sample. In the latter, two complexes could be distinguished, representing 1:1 and 1:2 RNA-to-ligand ratios, which are consistent with the crystallographic model showing two potential ANP77 binding sites (Figure 3C and 3D). The CD spectra of G2C4 RNA composed of five repeats did not indicate any major changes in the RNA structure upon ANP77 binding. Only additional ligand-binding-induced CD bands in the region of 300-380 nm were observed (Figure S2).

### X-ray structure versus biochemical data

The results of the biophysical evaluation can be interpreted in terms of structural data. In G2C4 RNA, ANP77 interacted with cytosine residues located in the single-stranded region. Although six H-bonds were formed between RNA and the ligand, the stacking interactions were limited, and the entropy of the cytosines was reduced, resulting in an elusive thermal effect. DSC measurements and mass spectra indicated that in solution, G2C4 existed as a tetramer folded into a triplex structure (Figure 3). The best condition for triplex formation was pH 6.0, at which two peaks were observed. The first peak likely corresponds to the melting of the Hoogsteen interactions in the base triples, whereas the second peak represents the melting of the slippery duplex. At pH 7.0, the protonation of cytosines required for base triple formation was more difficult, explaining the sole presence of duplex species. The DSC spectra at pH 5.3 also suggested the presence of one stable structure, presumably i-motif, which was easily formed under acidic conditions.^23^ Alternatively, stabilization of the triplex motif could occur, resulting in the merging of the duplex and tetramer signals into one peak.^19^

pH-dependent stabilization of the pyrimidine major groove triplexes was demonstrated by thermodynamic and NMR studies. Under acidic conditions, a higher rate of cytosine residue protonation results in the tightening of Hoogsteen interactions in triple base pairs.^24^ However, this requirement for protonation to form tertiary contacts does not explain the low stability of this motif in the cellular environment.^25^ Although the calculated pKa of the isolated cytosine was below pH 5.0, the structural context and solvent content imply the protonation of C, even under basic conditions.^26^ The pKa shift in cytosine has been observed in several crystal structures.^22, 27^ These studies demonstrated that formation of the C+GCA motif was crucial for proper folding and RNA activity. The assembly of G2C4 into the triple helix also re-quires the protonation of cytosines. Our crystallographic and biophysical results provided evidence that C+ species can be observed under near-physiological conditions, suggesting that they can also be stable in the cellular matrix. Similar conclusions were drawn in an NMR study of DNA containing G2C4 repeats.^10^

## CONCLUSIONS

Structural polymorphisms of the sense and antisense transcripts of HR result in complex pathological pathways in ALS/FTD. Our study demonstrates the potential of Crich G2C4 repeats to form higher-order structures. The “driving force” for RNA folding are cytosine residues. The limited nucleotide composition of G2C4 RNA results in a pKa shift of cytosines, extending their capabilities for Hbonding under near-physiological conditions. Consequently, a triplex structure containing C+•G-C and unique C•C+•C i-motif base triples is formed. The presented fold of G2C4 repeats is another example of RNA structural diversity and can be considered as a platform for RNA drug development against ALS/FTD.

ANP77 is a binding candidate for C-rich G2C4 RNA. It directly binds to adjacent cytosine residues and forms pseudo-canonical base pairs. This is our second study indicating that small molecules interact with the Watson-Crick edges of nucleotides. We previously showed that cyclic mismatch-binding ligand (CMBL) exhibited specificity toward adenosine residues in CAG repeats associated with polyglutamine disorders.^28^ We demonstrated that ANP77 and CMBL do not require the formation of “pockets” for specific RNA recognition but rather the presence of particular structural motifs. Bioinformatic tools are one strategy for design and identification of lead compounds.^14^ The crystallographic data for ANP77 and CMBL ligands and its derivatives can be used for in silico predictions, providing detailed information about their structure and interactions with RNA which could accelerate drug development against RNA-driven disorders.

## ASSOCIATED CONTENT

RNA synthesis, purification and crystallization, X-ray data collection, structure solution, and refinement, circular dichroism measurements, DSC measurements.

## Supporting Information

The Supporting Information is available free of charge at https://pubs.acs.org>

## AUTHOR INFORMATION

### Authors

**Leszek Błaszczyk** - Institute of Bioorganic Chemistry, Polish Academy of Sciences, Z. Noskowskiego 12/14, 61-704 Poznań, Poland

**Marcin Ryczek** - Institute of Bioorganic Chemistry, Polish Academy of Sciences, Z. Noskowskiego 12/14, 61-704 Poznań, Poland

**Bimolendu Das** - Department of Regulatory Bioorganic Chemistry, SANKEN (The Institute of Scientific and Industrial Research), Osaka University, 8-1 Mihogaoka, Ibaraki, Osaka 567-0047, Japan

**Martyna Mateja-Pluta** - Institute of Bioorganic Chemistry, Polish Academy of Sciences, Z. Noskowskiego 12/14, 61-704 Poznań, Poland

**Magdalena Bejger** - Institute of Bioorganic Chemistry, Polish Academy of Sciences, Z. Noskowskiego 12/14, 61-704 Poznań, Poland

**Kazuhiko Nakatani** - Department of Regulatory Bioorganic Chemistry, SANKEN (The Institute of Scientific and Industrial Research), Osaka University, 8-1 Mihogaoka, Ibaraki, Osaka 567-0047, Japan

## Author Contributions

### Notes

The authors declare no competing financial interest.

## ACKNOWLEDGMENT

This work was supported by National Science Centre grant UMO-2022/45/B/NZ7/03543. This work was partially supported by the JSPS KAKENHI Grant-in-Aid for Scientific Research (A) [22H00351] to K.N. For Open Access publishing, the authors have applied a CC-BY public copyright license to any Author Accepted Manuscript (AAM) version arising from this submission.

## TABLE OF CONTENTS

The first crystal structure of disease-related RNA CCCCGG in complex with synthetic ligand is reported. The ligand adopts a sandwich-like conformation and it mimics nucleobases by forming pseudo-canonical base pairs with cytosine residues of the triplex-like RNA. It demonstrates the structural polymorphism of cytosine-rich sequences and provide a template for further refinement of lead compounds for drug discovery against RNA-driven disorders.

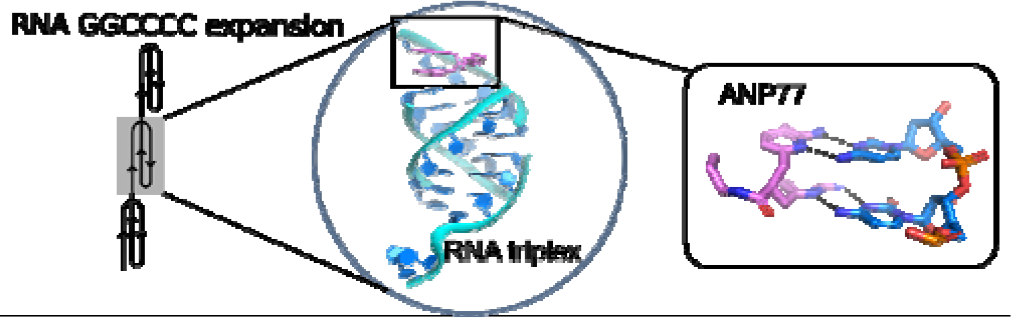

## REFERENCES

(1) Devenney, E. M.; Ahmed, R. M.; Hodges, J. R. Frontotemporal dementia. Handbook of clinical neurology 2019, 167, 279–299. DOI: 10.1016/B978-0-12-804766-8.00015-7. Geser, F.; Lee, V. M.; Trojanowski, J. Q. Amyotrophic lateral sclerosis and frontotemporal lobar degeneration: a spectrum of TDP-43 proteinopathies. Neuropathology : official journal of the Japanese Society of Neuropathology 2010, 30 (2), 103–112. DOI: 10.1111/j.1440-1789.2009.01091.x.

(2) DeJesus-Hernandez, M.; Mackenzie, I. R.; Boeve, B. F.; Boxer, A. L.; Baker, M.; Rutherford, N. J.; Nicholson, A. M.; Finch, N. A.; Flynn, H.; Adamson, J.; et al. Expanded GGGGCC hexanucleotide repeat in noncoding region of C9ORF72 causes chromosome 9p-linked FTD and ALS. Neuron 2011, 72 (2), 245–256. DOI: 10.1016/j.neuron.2011.09.011.

(3) Beck, J.; Poulter, M.; Hensman, D.; Rohrer, J. D.; Mahoney, C. J.; Adamson, G.; Campbell, T.; Uphill, J.; Borg, A.; Fratta, P.; et al. Large C9orf72 hexanucleotide repeat expansions are seen in multiple neurodegenerative syndromes and are more frequent than expected in the UK population. American journal of human genetics 2013, 92 (3), 345–353. DOI: 10.1016/j.ajhg.2013.01.011. van Blitterswijk, M.; DeJesus-Hernandez, M.; Niemantsverdriet, E.; Murray, M. E.; Heckman, M. G.; Diehl, N. N.; Brown, P. H.; Baker, M. C.; Finch, N. A.; Bauer, P. O.; et al. Association between repeat sizes and clinical and pathological characteristics in carriers of C9ORF72 repeat expansions (Xpansize-72): a cross-sectional cohort study. The Lancet. Neurology 2013, 12 (10), 978–988. DOI: 10.1016/S1474-4422(13)70210-2.

(4) Haeusler, A. R.; Donnelly, C. J.; Periz, G.; Simko, E. A.; Shaw, P. G.; Kim, M. S.; Maragakis, N. J.; Troncoso, J. C.; Pandey, A.; Sattler, R.; et al. C9orf72 nucleotide repeat structures initiate molecular cascades of disease. Nature 2014, 507 (7491), 195–200. DOI: 10.1038/nature13124.

(5) Kumar, V.; Kashav, T.; Islam, A.; Ahmad, F.; Hassan, M. I. Structural insight into C9orf72 hexanucleotide repeat expansions: Towards new therapeutic targets in FTD-ALS. Neurochemistry international 2016, 100, 11–20. DOI: 10.1016/j.neuint.2016.08.008.

(6) Gendron, T. F.; Bieniek, K. F.; Zhang, Y. J.; Jansen-West, K.; Ash, P. E.; Caulfield, T.; Daughrity, L.; Dunmore, J. H.; Castanedes-Casey, M.; Chew, J.; et al. Antisense transcripts of the expanded C9ORF72 hexanucleotide repeat form nuclear RNA foci and undergo repeat-associated non-ATG translation in c9FTD/ALS. Acta neuropathologica 2013, 126 (6), 829–844. DOI: 10.1007/s00401-013-1192-8. Mizielinska, S.; Lashley, T.; Norona, F. E.; Clayton, E. L.; Ridler, C. E.; Fratta, P.; Isaacs, A. M. C9orf72 frontotemporal lobar degeneration is characterised by frequent neuronal sense and antisense RNA foci. Acta neuropathologica 2013, 126 (6), 845–857. DOI: 10.1007/s00401-013-1200-z. Mori, K.; Lammich, S.; Mackenzie, I. R.; Forne, I.; Zilow, S.; Kretzschmar, H.; Edbauer, D.; Janssens, J.; Kleinberger, G.; Cruts, M.; et al. hnRNP A3 binds to GGGGCC repeats and is a constituent of p62-positive/TDP43-negative inclusions in the hippocampus of patients with C9orf72 mutations. Acta neuropathologica 2013, 125 (3), 413–423. DOI: 10.1007/s00401-013-1088-7.

(7) Mizielinska, S.; Gronke, S.; Niccoli, T.; Ridler, C. E.; Clayton, E. L.; Devoy, A.; Moens, T.; Norona, F. E.; Woollacott, I. O. C.; Pietrzyk, J.; et al. C9orf72 repeat expansions cause neurodegeneration in Drosophila through arginine-rich proteins. Science 2014, 345 (6201), 1192–1194. DOI: 10.1126/science.1256800. Mori, K.; Arzberger, T.; Grasser, F. A.; Gijselinck, I.; May, S.; Rentzsch, K.; Weng, S. M.; Schludi, M. H.; van der Zee, J.; Cruts, M.; et al. Bidirectional transcripts of the expanded C9orf72 hexanucleotide repeat are translated into aggregating dipeptide repeat proteins. Acta neuropathologica 2013, 126 (6), 881–893. DOI: 10.1007/s00401-013-1189-3. Mori, K.; Weng, S. M.; Arzberger, T.; May, S.; Rentzsch, K.; Kremmer, E.; Schmid, B.; Kretzschmar, H. A.; Cruts, M.; Van Broeckhoven, C.; et al. The C9orf72 GGGGCC repeat is translated into aggregating dipeptide-repeat proteins in FTLD/ALS. Science 2013, 339 (6125), 1335–1338. DOI: 10.1126/science.1232927.

(8) Bose, K.; Maity, A.; Ngo, K. H.; Vandana, J. J.; Shneider, N. A.; Phan, A. T. Formation of RNA G-wires by G(4)C(2) repeats associated with ALS and FTD. Biochemical and biophysical research communications 2022, 610, 113–118. DOI: 10.1016/j.bbrc.2022.03.162. Fratta, P.; Mizielinska, S.; Nicoll, A. J.; Zloh, M.; Fisher, E. M.; Parkinson, G.; Isaacs, A. M. C9orf72 hexanucleotide repeat associated with amyotrophic lateral sclerosis and frontotemporal dementia forms RNA G-quadruplexes. Scientific reports 2012, 2, 1016. DOI: 10.1038/srep01016. Jain, A.; Vale, R. D. RNA phase transitions in repeat expansion disorders. Nature 2017, 546 (7657), 243–247. DOI: 10.1038/nature22386.

(9) Su, Z.; Zhang, Y.; Gendron, T. F.; Bauer, P. O.; Chew, J.; Yang, W. Y.; Fostvedt, E.; Jansen-West, K.; Belzil, V. V.; Desaro, P.; et al. Discovery of a biomarker and lead small molecules to target r(GGGGCC)-associated defects in c9FTD/ALS. Neuron 2014, 83 (5), 1043–1050. DOI: 10.1016/j.neuron.2014.07.041.

(10) Kovanda, A.; Zalar, M.; Sket, P.; Plavec, J.; Rogelj, B. Anti-sense DNA d(GGCCCC)n expansions in C9ORF72 form i-motifs and protonated hairpins. Scientific reports 2015, 5, 17944. DOI: 10.1038/srep17944.

(11) Dodd, D. W.; Tomchick, D. R.; Corey, D. R.; Gagnon, K. T. Pathogenic C9ORF72 Antisense Repeat RNA Forms a Double Helix with Tandem C:C Mismatches. Biochemistry 2016, 55 (9), 1283–1286. DOI: 10.1021/acs.biochem.6b00136.

(12) Simone, R.; Balendra, R.; Moens, T. G.; Preza, E.; Wilson, K. M.; Heslegrave, A.; Woodling, N. S.; Niccoli, T.; Gilbert-Jaramillo, J.; Abdelkarim, S.; et al. G-quadruplex-binding small molecules ameliorate C9orf72 FTD/ALS pathology in vitro and in vivo. EMBO molecular medicine 2018, 10 (1), 22–31. DOI: 10.15252/emmm.201707850. Ursu, A.; Baisden, J. T.; Bush, J. A.; Taghavi, A.; Choudhary, S.; Zhang, Y. J.; Gendron, T. F.; Petrucelli, L.; Yildirim, I.; Disney, M. D. A Small Molecule Exploits Hidden Structural Features within the RNA Repeat Expansion That Causes c9ALS/FTD and Rescues Pathological Hallmarks. ACS chemical neuroscience 2021, 12 (21), 4076–4089. DOI: 10.1021/acschemneuro.1c00470. Ursu, A.; Wang, K. W.; Bush, J. A.; Choudhary, S.; Chen, J. L.; Baisden, J. T.; Zhang, Y. J.; Gendron, T. F.; Petrucelli, L.; Yildirim, I.; et al. Structural Features of Small Molecules Targeting the RNA Repeat Expansion That Causes Genetically Defined ALS/FTD. ACS chemical biology 2020, 15 (12), 3112–3123. DOI: 10.1021/acschembio.0c00049.

(13) Wang, Z. F.; Ursu, A.; Childs-Disney, J. L.; Guertler, R.; Yang, W. Y.; Bernat, V.; Rzuczek, S. G.; Fuerst, R.; Zhang, Y. J.; Gendron, T. F.; et al. The Hairpin Form of r(G(4)C(2))(exp) in c9ALS/FTD Is Repeat-Associated Non-ATG Translated and a Target for Bioactive Small Molecules. Cell chemical biology 2019, 26 (2), 179–190 e112. DOI: 10.1016/j.chembiol.2018.10.018.

(14) Childs-Disney, J. L.; Yang, X.; Gibaut, Q. M. R.; Tong, Y.; Batey, R. T.; Disney, M. D. Targeting RNA structures with small molecules. Nature reviews. Drug discovery 2022, 21 (10), 736–762. DOI: 10.1038/s41573-022-00521-4.

(15) Warner, K. D.; Hajdin, C. E.; Weeks, K. M. Principles for targeting RNA with drug-like small molecules. Nature reviews. Drug discovery 2018, 17 (8), 547–558. DOI: 10.1038/nrd.2018.93.

(16) Das, B.; Murata, A.; Nakatani, K. A small-molecule fluorescence probe ANP77 for sensing RNA internal loop of C, U and A/CC motifs and their binding molecules. Nucleic acids research 2021, 49 (15), 8462–8470. DOI: 10.1093/nar/gkab650.

(17) Das, B.; Nagano, K.; Kawai, G.; Murata, A.; Nakatani, K. 2-Amino-1,8-naphthyridine Dimer (ANP77), a High-Affinity Binder to the Internal Loops of C/CC and T/CC Sites in Double-Stranded DNA. The Journal of organic chemistry 2022, 87 (1), 340–350. DOI: 10.1021/acs.joc.1c02383.

(18) Brown, J. A. Unraveling the structure and biological functions of RNA triple helices. Wiley interdisciplinary reviews. RNA 2020, 11 (6), e1598. DOI: 10.1002/wrna.1598.

(19) Devi, G.; Zhou, Y.; Zhong, Z.; Toh, D. F.; Chen, G. RNA triplexes: from structural principles to biological and biotech applications. Wiley interdisciplinary reviews. RNA 2015, 6 (1), 111–128. DOI: 10.1002/wrna.1261.

(20) Kiliszek, A.; Kierzek, R.; Krzyzosiak, W. J.; Rypniewski, W. Crystallographic characterization of CCG repeats. Nucleic acids research 2012, 40 (16), 8155–8162. DOI: 10.1093/nar/gks557. Rypniewski, W.; Banaszak, K.; Kulinski, T.; Kiliszek, A. Watson-Crick-like pairs in CCUG repeats: evidence for tautomeric shifts or protonation. Rna 2016, 22 (1), 22–31. DOI: 10.1261/rna.052399.115.

(21) Brown, J. A.; Bulkley, D.; Wang, J.; Valenstein, M. L.; Yario, T. A.; Steitz, T. A.; Steitz, J. A. Structural insights into the stabilization of MALAT1 noncoding RNA by a bipartite triple helix. Nature structural & molecular biology 2014, 21 (7), 633–640. DOI: 10.1038/nsmb.2844. Brown, J. A.; Valenstein, M. L.; Yario, T. A.; Tycowski, K. T.; Steitz, J. A. Formation of triple-helical structures by the 3’-end sequences of MALAT1 and MENbeta noncoding RNAs. Proceedings of the National Academy of Sciences of the United States of America 2012, 109 (47), 19202–19207. DOI: 10.1073/pnas.1217338109. Chen, G.; Chang, K. Y.; Chou, M. Y.; Bustamante, C.; Tinoco, I., Jr. Triplex structures in an RNA pseudoknot enhance mechanical stability and increase efficiency of -1 ribosomal frameshifting. Proceedings of the National Academy of Sciences of the United States of America 2009, 106 (31), 12706–12711. DOI: 10.1073/pnas.0905046106. Gilbert, S. D.; Rambo, R. P.; Van Tyne, D.; Batey, R. T. Structure of the SAM-II riboswitch bound to S-adenosylmethionine. Nature structural & molecular biology 2008, 15 (2), 177–182. DOI: 10.1038/nsmb.1371. Klein, D. J.; Ferre-D’Amare, A. R. Structural basis of glmS ribozyme activation by glucosamine-6-phosphate. Science 2006, 313 (5794), 1752–1756. DOI: 10.1126/science.1129666. Liberman, J. A.; Salim, M.; Krucinska, J.; Wedekind, J. E. Structure of a class II preQ1 riboswitch reveals ligand recognition by a new fold. Nature chemical biology 2013, 9 (6), 353–355. DOI: 10.1038/nchembio.1231. Qiao, F.; Cech, T. R. Triple-helix structure in telomerase RNA contributes to catalysis. Nature structural & molecular biology 2008, 15 (6), 634–640. DOI: 10.1038/nsmb.1420. Shefer, K.; Brown, Y.; Gorkovoy, V.; Nussbaum, T.; Ulyanov, N. B.; Tzfati, Y. A triple helix within a pseudoknot is a conserved and essential element of telomerase RNA. Molecular and cellular biology 2007, 27 (6), 2130–2143. DOI: 10.1128/MCB.01826-06. Theimer, C. A.; Blois, C. A.; Feigon, J. Structure of the human telomerase RNA pseudoknot reveals conserved tertiary interactions essential for function. Molecular cell 2005, 17 (5), 671–682. DOI: 10.1016/j.molcel.2005.01.017. Ulyanov, N. B.; Shefer, K.; James, T. L.; Tzfati, Y. Pseudoknot structures with conserved base triples in telomerase RNAs of ciliates. Nucleic acids research 2007, 35 (18), 6150–6160. DOI: 10.1093/nar/gkm660.

(22) Nixon, P. L.; Rangan, A.; Kim, Y. G.; Rich, A.; Hoffman, D. W.; Hennig, M.; Giedroc, D. P. Solution structure of a luteoviral P1-P2 frameshifting mRNA pseudoknot. Journal of molecular biology 2002, 322 (3), 621–633. DOI: 10.1016/s0022-2836(02)00779-9.

(23) Benabou, S.; Avino, A.; Eritja, R.; Gonzalez, C.; Gargallo, R. Fundamental aspects of the nucleic acid i-motif structures. Rsc Adv 2014, 4 (51), 26956–26980. DOI: 10.1039/c4ra02129k. Snoussi, K.; Nonin-Lecomte, S.; Leroy, J. L. The RNA i-motif. Journal of molecular biology 2001, 309 (1), 139–153. DOI: 10.1006/jmbi.2001.4618.

(24) Geierstanger, B. H.; Wemmer, D. E. Complexes of the Minor-Groove of DNA. Annu Rev Bioph Biom 1995, 24, 463–493. DOI: DOI 10.1146/annurev.bb.24.060195.002335. Holland, J. A.; Hoffman, D. W. Structural features and stability of an RNA triple helix in solution. Nucleic acids research 1996, 24 (14), 2841–2848. DOI: 10.1093/nar/24.14.2841.

(25) Moody, E. M.; Lecomte, J. T.; Bevilacqua, P. C. Linkage between proton binding and folding in RNA: a thermodynamic framework and its experimental application for investigating pKa shifting. Rna 2005, 11 (2), 157–172. DOI: 10.1261/rna.7177505.

(26) Jones, E. L.; Mlotkowski, A. J.; Hebert, S. P.; Schlegel, H. B.; Chow, C. S. Calculations of pK(a) Values for a Series of Naturally Occurring Modified Nucleobases. The journal of physical chemistry. A 2022, 126 (9), 1518–1529. DOI: 10.1021/acs.jpca.1c10905. Tang, C. L.; Alexov, E.; Pyle, A. M.; Honig, B. Calculation of pKas in RNA: on the structural origins and functional roles of protonated nucleotides. Journal of molecular biology 2007, 366 (5), 1475–1496. DOI: 10.1016/j.jmb.2006.12.001.

(27) Ferre-D’Amare, A. R.; Zhou, K.; Doudna, J. A. Crystal structure of a hepatitis delta virus ribozyme. Nature 1998, 395 (6702), 567–574. DOI: 10.1038/26912. Ke, A.; Zhou, K.; Ding, F.; Cate, J. H.; Doudna, J. A. A conformational switch controls hepatitis delta virus ribozyme catalysis. Nature 2004, 429 (6988), 201–205. DOI: 10.1038/nature02522. Nixon, P. L.; Cornish, P. V.; Suram, S. V.; Giedroc, D. P. Thermodynamic analysis of conserved loop-stem interactions in P1-P2 frameshifting RNA pseudoknots from plant Luteoviridae. Biochemistry 2002, 41 (34), 10665–10674. DOI: 10.1021/bi025843c. Nixon, P. L.; Giedroc, D. P. Energetics of a strongly pH dependent RNA tertiary structure in a frameshifting pseudoknot. Journal of molecular biology 2000, 296 (2), 659–671. DOI: 10.1006/jmbi.1999.3464. Su, L.; Chen, L.; Egli, M.; Berger, J. M.; Rich, A. Minor groove RNA triplex in the crystal structure of a ribosomal frameshifting viral pseudoknot. Nature structural biology 1999, 6 (3), 285–292. DOI: 10.1038/6722.

(28) Mukherjee, S.; Blaszczyk, L.; Rypniewski, W.; Falschlunger, C.; Micura, R.; Murata, A.; Dohno, C.; Nakatani, K.; Kiliszek, A. Structural insights into synthetic ligands targeting A-A pairs in disease-related CAG RNA repeats. Nucleic acids research 2019, 47 (20), 10906–10913. DOI: 10.1093/nar/gkz832.

