## Supplemental Information for "Antisense RNA C9orf72 Hexanucleotide Repeat Associated With Amyotrophic Lateral Sclerosis and Frontotemporal Dementia Forms A Triplex-Like Structure and Binds Small Synthetic Ligand"

**Abstract:** The abnormal expansion of GGGGCC/CCCCGG hexanucleotide repeats (HR) in *C9orf72* is associated with familial amyotrophic lateral sclerosis (ALS) and frontotemporal dementia (FTD). Structural polymorphisms of HR result in the multifactorial pathomechanism of ALS/FTD. Consequently, many ongoing studies are focused at developing therapies targeting pathogenic HR RNA. One of them involves small molecules blocking the sequestration of important proteins, preventing the formation of toxic nuclear foci. However, rational design of potential therapeutics is hindered by limited number of structural studies of RNA-ligand complexes. We determined the crystal structure of antisense HR RNA in complex with ANP77 ligand and in the free form. HR RNA folds into a triplex structure composed of four RNA chains. ANP77 interacted with two neighboring single-stranded cytosines to form pseudo-canonical base pairs by adopting sandwich-like conformation and adjusting the position of its naphthyridine units to the helical twist of the RNA. In the unliganded structure, the cytosines formed a peculiar triplex i-motif, assembled by *trans* C•C$^+$ pair and a third cytosine located at the Hoogsteen edge of the C•C$^+$ pair. These results extend our knowledge of the structural polymorphisms of HR and can be used for the rational design of small molecules targeting disease-related RNAs.

### Experimental Procedures

#### Synthesis, purification and crystallization of RNA oligomers

The oligomer was synthesized by the solid phase method using Applied Biosystems DNA/RNA synthesizer and TOM protected phosphoramidites. The synthesis was carried out in DMT-ON mode and RNA was purified according to the protocol suitable for Glen-Pak cartridges (Glen Research). The purified RNA was lyophilized under vacuum using a Speed-Vac and stored at -20°C. Before crystallization the RNA was dissolved in buffer containing 10 mM sodium cacodylate pH 7.0 and 100 mM NaCl. The final concentration of RNA was 0.5 mM. The oligomer was denatured for 5 min at 95°C and snap-cooled on ice for 10 min. Then RNA sample was incubated for 20 min in RT with or without 0.75 mM ANP77. Crystals were grown by the sitting drop method at 19°C. The crystals of RNA-ligand complex grew in 80 mM NaCl, 40 mM Na cacodylate trihydrate pH 6.0, 45% v/v MPD, 12 mM spermine tetrahydrochloride, while unliganded RNA oligomer was crystallized in 80 mM NaCl, 20 mM magnesium chloride hexahydrate, 40 mM Na cacodylate trihydrate pH 6.0, 35% v/v MPD and 12 mM spermine tetrahydrochloride.

#### X-ray data collection, structure solution, and refinement

X-ray diffraction data were collected on in house diffractometer XtaLab Synergy-R Rigaku. The data were integrated using CrysAlisPro software provided by Rigaku. The data were scaled using SCALA from CCP4 program suite.[1] The structure of RNA-ligand complex was solved by molecular replacement with PHASER using part of RNA model containing $C_4G_2$ repeats (PDB code: 5ew7).[2] The phases of unliganded RNA model were taken from RNA-ligand structure. Early stages of the refinement were done using the Refmac5 from the CCP4 program suite. Further refinement was carried out with PHENIX.[3] The manual model building was done using Coot.[4] Restraints for the ANP77 ligand was generated by the grade web server developed by Global Phasing (http://grade.globalphasing.org). All pictures were drawn using PyMOL.[5] Atomic coordinates of the crystallographic models have been deposited with the Protein Data Bank (accession codes: 8QMH, 8QMI).The statistics of X-ray data collection and refinement are summarized in Table S2. X-ray diffraction images are deposited in the MX-RDR database (Macromolecular Xtallography Raw Data Repository) (https://mxrdr.icm.edu.pl/) (doi:10.18150/AJNJBE for RNA-ANP77 structure and doi:10.18150/FESTPM for native RNA model). X-ray data and model refinement statistics are summarized in Table S1.

#### CSI-TOF-MS measurements

Time-of-flight mass spectrometry analysis by cold spray ionization (CSI-TOF MS) data were obtained in negative mode using JEOL JMS-T100LP AccuTOF LC-plus 4G mass spectrometer. Samples containing the $G_2C_4$ RNA (20 μM) with or without ANP77 (50 μM) in an 8:2 mixture of water and methanol containing ammonium acetate (250 mM) were sprayed at a flow rate of 14μL $min^{-1}$. During the injection, the spray temperature was set at −10°C.

#### Circular Dichroism measurements

Circular dichroism (CD) experiments were carried out on a J-725 CD spectrometer (JASCO) using a 10 mm path length cell at room temperature. The CD spectra of repeat $G_2C_4$ RNA (2.5 μM) were measured with sodium cacodylate buffer (10 mM, pH 7.0) containing 100 mM NaCl in the absence and presence of ANP77 (25 μM).

**DSC measurements**

The RNA oligomers were dissolved in 100 mM sodium chloride and 10 mM sodium cacodylate buffer adjusted to pH 7.0, 6.0, or 5.2. The final concentration of RNA was 300 μM. Samples were dialyzed against buffer overnight at 4°C to equilibrate RNA and the reference solutions. Before DSC measurements, RNA samples were denatured at 95°C for 5 min and snap-cooled on ice for 10 min. Next, the RNA was renatured for 20 min in RT with or without 300 μM ANP77 ligand. DSC experiments were performed on a MicroCal PEAQ-DSC calorimeter (Malvern Instruments Ltd.) Each measurement was carried out in five cycles of heating and cooling in the range of 2–110°C and a scan rate of 1°C/min. First, reference scans of buffer were performed to establish the instrument thermal history and to reach a near perfect baseline repeatability. The results were analyzed using dedicated software implemented by Malvern Instruments. The melting temperature (Tm) was calculated by applying a two-state model fitting.

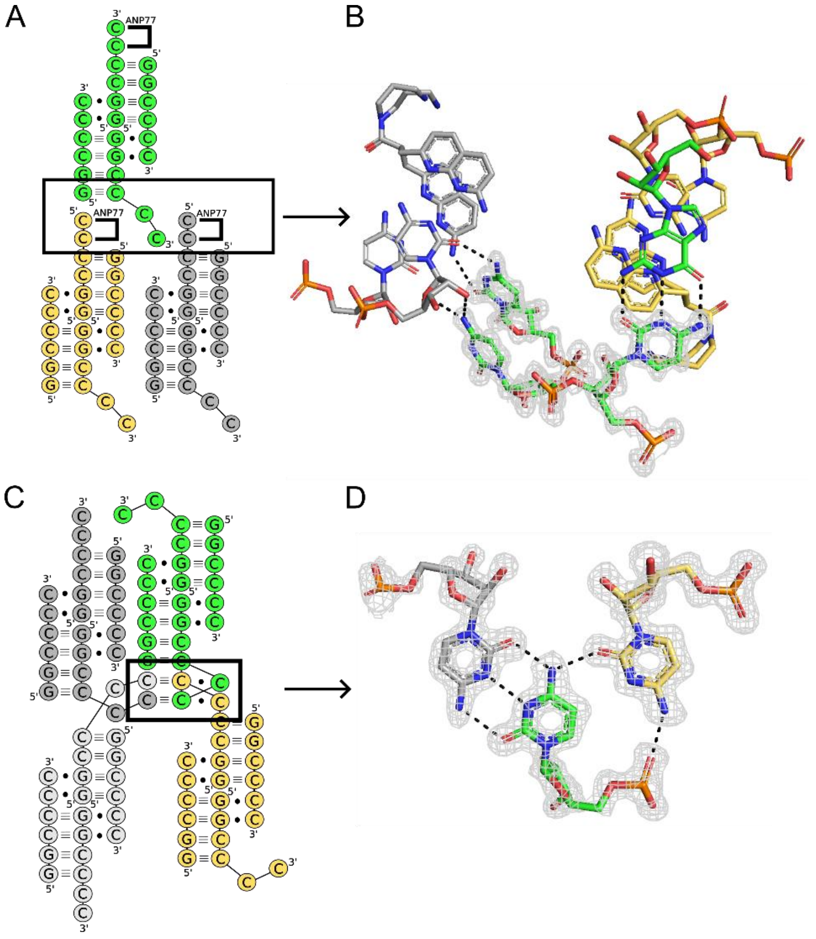


**Figure S1.** Crystal contacts observed in the crystal lattice of RNA-ligand complex (A-B) and in uliganded model (C-D). (A and C) the schematic representation of crystal contacts between symmetry-related molecules of $G_2C_4$ RNA tetramers. In the liganded structure (A) the overhanging cytosines are located in the vicinity of pseudo-canonical base pairs formed between ANP77 and cytosines of symmetry-related RNA tetramer (black box). (B) Details of H-bond pattern observed between overhanging cytosines (green sticks) and cytosines and ANP77 (grey sticks) from symmetry-related molecule. (C-D) In the unliganded structure the respective cytosine residues are involved in formation of two unique base triples (black box).The 2Fo–Fc electron density map (gray) is contoured at the 1σ level. The H-bonds are denoted as black dashed lines.

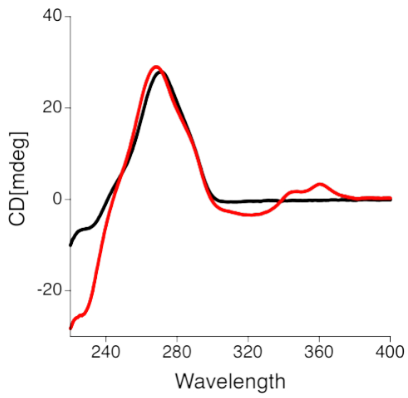


**Figure S2.** CD spectra of $G_2C_4$ RNA five repeats in the absence (black line) and presence (red line) of ANP77 ligand. Concentrations of RNA and ANP77 were 2.5 and 25 μM, respectively.

**Table S1.** Summary of X-ray data and model refinement statistics.

| | $G_2C_4$ + ANP77 | $G_2C_4$ native |
|---|---|---|
| **Data Collection** | | |
| X-ray source | XtaLAB Synergy-R, HyPix | XtaLAB Synergy-R, HyPix |
| Space group | $C222_1$ | $C222_1$ |
| Cell parameters (Å) | a = 29.34, b = 45.18, c = 86.28 | a = 41.76, b = 46.64, c = 51.49 |
| Resolution (Å) | 21.85-1.10 (1.16-1.1)[a] | 21.25-1.50 (1.58-1.50)# |
| $R_{merge}$ | 0.076 (0.560) | 0.116 (0.553) |
| I/σ | 26.9 (4.4) | 10.0 (2.4) |
| $CC_{1/2}$ | 0.99 (0.89) | 0.99 (0.86) |
| Completeness (%) | 99.9 (99.4) | 99.7 (100) |
| Redundancy | 16 (11.0) | 7.3 (4.8) |
| Number of unique reflections | 23771 | 8334 |
| **Refinement** | | |
| Software | Refmac 5.8.0405/ Phenix (1.20.1) | |
| Number of reflections: work/test | 22774/479 | 7905/404 |
| Overall mean B value (Å$^2$) | 13.03 | 8.62 |
| $R_{work}/R_{free}$ | 0.1218/0.1737 | 0.1680/0.2254 |
| RNA atoms | 724 | 555 |
| Water molecules | 108 | 108 |
| Ligand molecules | 1 ANP77 | 2 $Mg^{2+}$ ions |
| RMSD in bonds (Å) | 0.008 | 0.007 |
| RMSD in angles (°) | 1.4 | 1.9 |
| PDB code | 8QMH | 8QMI |
| x-ray images | doi:10.18150/AJNJBE | doi:10.18150/FESTPM |

[a]Values in parentheses are for the last resolution shell

**Table S2.** Summary of Tm melting temperature measured by DSC in different pH.

| | $Tm_1$ (°C) | $Tm/Tm_2$ (°C) |
|---|---|---|
| pH 5.26 native | | 60.04 |
| pH 5.26 + ANP77 | | 61.29 |
| pH 6.0 native | 37.42 | 57.24 |
| pH 6.0 + ANP77 | 41.95 | 57.90 |
| pH 7.0 native | | 55.92 |
| pH 7.0 + ANP77 | | 54.92 |

### References

(1) Agirre, J.; Atanasova, M.; Bagdonas, H.; Ballard, C. B.; Basle, A.; Beilsten-Edmands, J.; Borges, R. J.; Brown, D. G.; Burgos-Marmol, J. J.; Berrisford, J. M.; et al. The CCP4 suite: integrative software for macromolecular crystallography. *Acta crystallographica. Section D, Structural biology* **2023**, *79* (Pt 6), 449-461. DOI: 10.1107/S2059798323003595. Evans, P. Scaling and assessment of data quality. *Acta crystallographica. Section D, Biological crystallography* **2006**, *62* (Pt 1), 72-82. DOI: 10.1107/S0907444905036693.

(2) McCoy, A. J.; Grosse-Kunstleve, R. W.; Adams, P. D.; Winn, M. D.; Storoni, L. C.; Read, R. J. Phaser crystallographic software. *Journal of applied crystallography* **2007**, *40* (Pt 4), 658-674. DOI: 10.1107/S0021889807021206.

(3) Afonine, P. V.; Grosse-Kunstleve, R. W.; Echols, N.; Headd, J. J.; Moriarty, N. W.; Mustyakimov, M.; Terwilliger, T. C.; Urzhumtsev, A.; Zwart, P. H.; Adams, P. D. Towards automated crystallographic structure refinement with phenix.refine. *Acta crystallographica. Section D, Biological crystallography* **2012**, *68* (Pt 4), 352-367. DOI: 10.1107/S0907444912001308. Kovalevskiy, O.; Nicholls, R. A.; Long, F.; Carlon, A.; Murshudov, G. N. Overview of refinement procedures within REFMAC5: utilizing data from different sources. *Acta crystallographica. Section D, Structural biology* **2018**, *74* (Pt 3), 215-227. DOI: 10.1107/S2059798318000979.

(4) Emsley, P.; Lohkamp, B.; Scott, W. G.; Cowtan, K. Features and development of Coot. *Acta crystallographica. Section D, Biological crystallography* **2010**, *66* (Pt 4), 486-501. DOI: 10.1107/S0907444910007493.

(5) The PyMOL Molecular Graphics System, Version 1.2r3pre, Schrödinger, LLC.
